# Intrinsic Organization of Contrast Sensitivity in Human Vision

**DOI:** 10.64898/2026.09.07.749815

**Authors:** Alexander Minns, Sergei Gepshtein, Natalia Janson, Sergey Savel’ev

## Abstract

Natural vision operates over an enormous range of luminance while preserving sensitivity to both fine and coarse spatial structure. As luminance increases, the peak of the contrast sensitivity function shifts toward higher spatial frequencies. Existing accounts explain how luminance rescales the amplitude of neural responses through gain control, but not why preferred spatial frequency should reorganize systematically with luminance. Here we propose that recurrent excitatory–inhibitory interactions generate an intrinsic spatial frequency, set by the coupling structure of the circuit, that shapes the network’s resonance and thereby predicts changes in preferred spatial frequency. Changes in mean luminance shift this intrinsic spatial frequency by altering the effective balance between excitation and inhibition, thereby moving the network between dynamical regimes rather than simply rescaling its responses. To test this framework, we measured human contrast sensitivity across a finely sampled range of luminance. Preferred spatial frequency followed the predicted course: approximately constant at low luminance, then rising sharply over a narrow range in an S-shaped transition. These results support a circuit-based account in which changes in cortical state reorganize spatial selectivity, linking luminance-dependent perception to the dynamics of cortical computation.

**Significance Statement:** The visual system maintains sensitivity across enormous changes in environmental illumination, yet the neural mechanism underlying this adaptability remains unclear. Existing theories explain how luminance changes response gain but not why the preferred spatial scale of visual processing changes with luminance. By combining psychophysical measurements with a recurrent excitatory–inhibitory network model, we show that preferred spatial frequency changes systematically with luminance and that these changes can arise from shifts in the intrinsic spatial frequency of the circuit. These findings provide a mechanistic link between recurrent cortical dynamics and luminance-dependent spatial tuning, suggesting that spatial-frequency selectivity can emerge from the state of the recurrent network.

## Introduction

How the visual system preserves selective sensitivity across the enormous range of natural luminance has long interested vision scientists [Kelly, 1975, Watson and Ahumada, 2005, Zemon et al., 2023]. This presents a fundamental computational challenge: changes in luminance alter both the amount and reliability of visual information reaching the retina, yet the visual system must continue to encode behaviourally relevant spatial information across these changing conditions. Consequently, observers are able to discern both fine and coarse spatial structure over a wide range of luminance levels, and the contrast sensitivity function (CSF) retains its characteristic form [Van Nes et al., 1967, Kelly, 1972, De Valois et al., 1974].

Early work on how visual sensitivity changes with luminance sought general principles to capture that dependence. Measurements of the CSF showed that its peak shifts toward higher spatial frequencies as luminance increases [Van Nes et al., 1967, Kelly, 1972], a pattern commonly explained in terms of efficiency: limits imposed by photon noise, neural noise, and the spatial scale over which visual signals can be reliably encoded [Mazade et al., 2022, Ghodrati et al., 2019]. Within this view, low luminance reduces the signal-to-noise ratio, making reliable detection of fine spatial structure more difficult and favoring integration over larger spatial scales, whereas increasing luminance progressively permits finer spatial detail to be resolved. Although these studies established that contrast sensitivity increases with luminance and that its peak shifts accordingly, the comparatively coarse sampling of luminance provided limited insight into the circuit mechanisms underlying the reorganization of spatial selectivity.

Subsequent work sought physiological explanations for how sensitivity changes with luminance. Many influential models framed these effects in terms of gain control [Carandini and Heeger, 2012, Ohzawa et al., 1985], typically formalized through divisive normalization, whereby a neuron’s response is scaled by the activity of a broader neural population [Heeger, 1992]. In such models, luminance modulates the gain applied to neural activity, thereby adjusting the effective contrast that neurons represent [Cox et al., 1999, De Valois et al., 1982]. While these approaches capture important aspects of sensitivity regulation, they primarily describe how luminance-dependent changes in neural activity are transformed into changes in response gain. However, they leave unresolved how changes in luminance give rise to changes in preferred spatial frequency, or how distinct tuning regimes might emerge from the underlying circuit dynamics.

Here we examine whether recurrent excitatory–inhibitory dynamics provide a mechanistic framework for understanding how the contrast sensitivity function adapts to changes in luminance. In recurrent excitatory–inhibitory networks, spatial interactions can give rise to preferred modes of activity characterized by an intrinsic spatial frequency, determined by the coupling structure of the circuit [Wilson and Cowan, 1972, 1973, Gepshtein et al., 2022]. Recent theoretical work has shown that such intrinsic spatial frequencies emerge naturally from recurrent cortical dynamics [Davis et al., 2024]. This intrinsic spatial frequency shapes the spatial resonance of the circuit and thereby the preferred spatial frequency observed in the CSF. Within this framework, changes in mean luminance modulate the background activity of the network, thereby modifying effective recurrent coupling and, in turn, altering the intrinsic spatial frequency of the system. If preferred spatial frequency is linked to the intrinsic spatial frequency of the recurrent circuit, then luminance-dependent changes in cortical state should produce corresponding changes in perceptual tuning.

Guided by these predictions, we measured human contrast sensitivity functions across a finely sampled luminance range and identified the spatial frequency at which sensitivity peaked. We then compared these measurements with the predictions of the recurrent excitatory–inhibitory framework to test whether the observed changes in preferred spatial frequency are consistent with luminance-dependent changes in effective excitatory–inhibitory balance and the resulting shift in the circuit’s intrinsic spatial frequency.

### Model Setup

A common approach to modelling neural circuits with interacting excitatory and inhibitory populations is the framework introduced by Wilson and Cowan [Wilson and Cowan, 1972, 1973] (Fig. 1A–B). This model provides a concise description of how local excitatory and inhibitory activity evolves over time and has been used to account for a wide range of experimental findings. In its simplest interpretation, excitatory activity tends to increase the activity of both excitatory and inhibitory neurons through recurrent and lateral connections, while inhibitory activity suppresses the activity of both populations. Mathematically, the Wilson–Cowan model is expressed as a system of coupled ordinary differential equations. This formalism captures many aspects of population dynamics, yet it is less suited to describing spatially extended phenomena such as traveling or standing waves, which arise naturally in distributed networks. Recent work [Gepshtein et al., 2022, Cruddas et al., 2026] has shown that the Wilson–Cowan equations can be reformulated as nonlinear partial differential equations. This reformulation treats neural tissue as a continuous medium when activity varies smoothly across space, and it provides a convenient way to study wave-like behavior such as propagation, interference, and spatial pattern formation. In contrast to neural field theory, which typically employs wave equations analogous to those used in field theory [Cruddas et al., 2026], the wave equations derived from the Wilson–Cowan model [Gepshtein et al., 2022] differ in that they are grounded in coupled excitatory and inhibitory population dynamics and thus directly reflect the organization of neural tissue.

**Figure 1:**
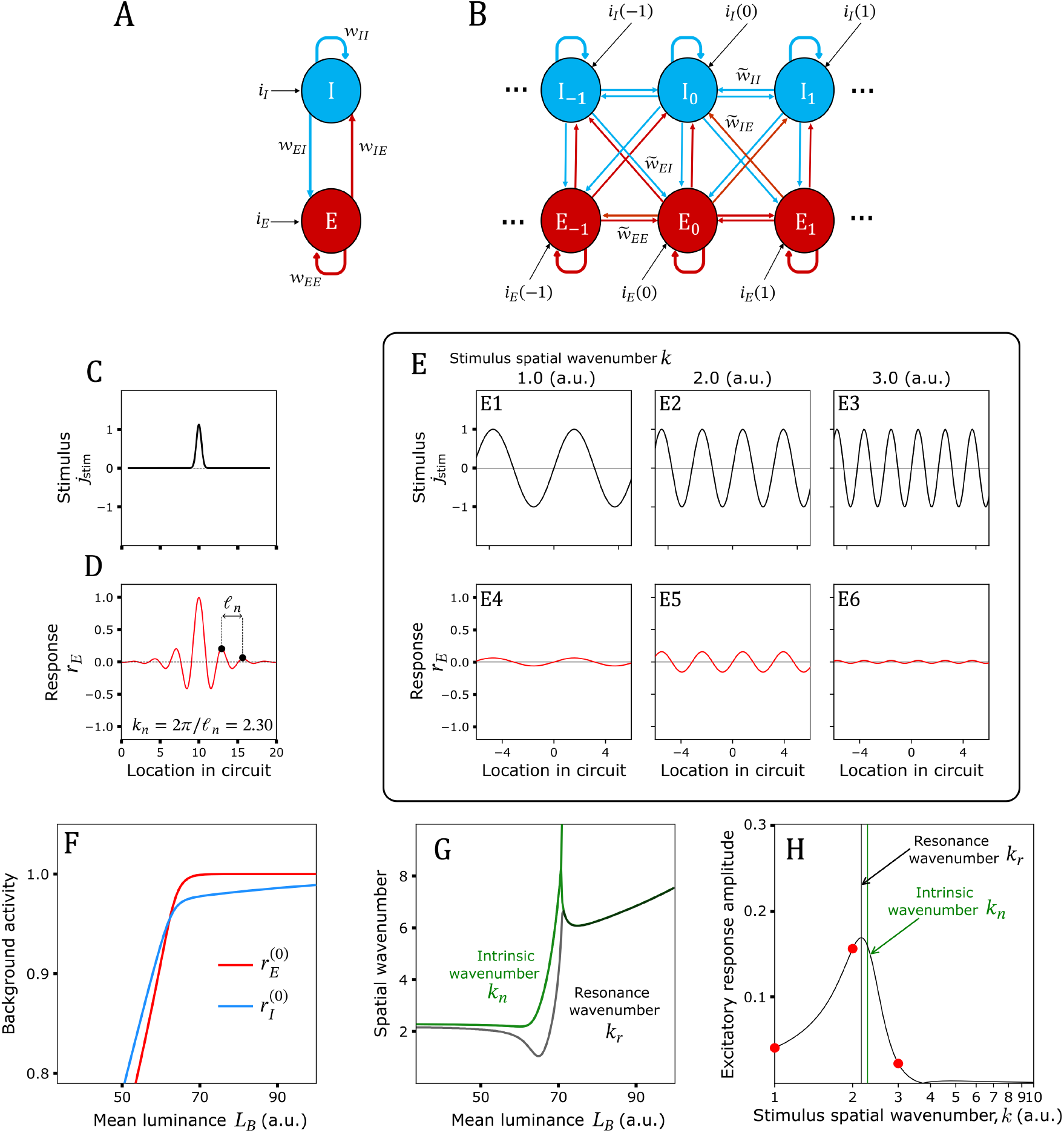
Model of excitatory–inhibitory network and its selectivity. (**A**) Wilson–Cowan motif with interacting excitatory (E) and inhibitory (I) populations, local coupling weights w_*PQ*_, and partitioned external input, where *P, Q ∈* {*E, I*}. (**B**) Spatially coupled chain of motifs. Here *i*_*E*_(*l*) and *i*_*I*_(*l*) are the external inputs at location *l*, and 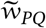 are the lateral coupling weights. (**C–D**) Localized input and corresponding excitatory Green’s-function response at *L*_*B*_ = 0. The response oscillates with intrinsic wavelength *l*_*n*_, where *k*_*n*_ = 2*π*/*l*_*n*_, and its envelope decays at rate *λ*. (**E**) Periodic stimuli of increasing spatial wavenumber *k* (E1–E3) and corresponding excitatory Green’s-function responses at *L*_*B*_ = 0 (E4–E6), with maximal amplification near the intrinsic wavenumber *k*_*n*_. (**F**) Baseline excitatory and inhibitory activities as functions of mean luminance *L*_*B*_, showing nonlinear saturation and a transition from *r*_*I*_ > *r*_*E*_ to *r*_*E*_ > *r*_*I*_. (**G**) Two real-part branches of the intrinsic wavenumber *k*_*n*_(*L*_*B*_) (green) and the resonance wavenumber *k*_*r*_(*L*_*B*_) (black). At each luminance, *k*_*r*_ is obtained from the maximum of the excitatory response coefficient *ξ(k, L*_*B*_). (**H**) Positive excitatory response amplitude obtained from the *L*_*B*_ = 0 Green’s-function calculation, plotted as a function of stimulus spatial wavenumber *k*. The response peaks at the resonance wavenumber *k*_*r*_, which lies near the intrinsic wavenumber *k*_*n*_. Red circles identify the wavenumbers used in E4–E6.

In our formalism, neural activity has the form of waves propagating in a network of coupled excitatory and inhibitory neurons (Fig. 1A and Materials and Methods). Activity spreads across neighboring locations, allowing spatial patterns of activation to propagate and interact (Fig. 1C–D). The strength of these interactions is governed by a set of coupling parameters (“weights”) that determine how activity is exchanged within and between the excitatory and inhibitory populations, both locally and across space. Together, these interactions define the spatial dynamics in neural space and give rise to wave-like responses [Gepshtein et al., 2022].

We consider the response of the network to a luminance grating, characterized by its spatial frequency and amplitude, presented on top of a uniform background luminance. The background luminance sets the baseline level of activity in both populations in the absence of structured input.

To understand how background luminance affects spatial selectivity, it is useful to separate the static, spatially homogeneous baseline activity from the stimulus-driven responses. The baseline corresponds to the operating state of the network under uniform input, whereas the stimulus introduces spatially varying perturbations around this state (see Materials and Methods). A key consequence of this separation is that the baseline activity alters how the network responds to the spatial structure of the stimulus. Because neural responses are nonlinear, changes in baseline activity modify the effective strength of interactions within the circuit. In other words, luminance shifts the operating point of the system and thereby changes the effective coupling between neurons (i.e., renormalizes it), without changing the underlying connectivity.

This renormalization affects both local interactions and spatial spread, leading to systematic changes in how activity propagates across the network. As a result, the circuit exhibits an intrinsic spatial frequency, determined by its effective coupling parameters, at which activity is preferentially amplified. Because effective coupling depends on baseline activity, and baseline activity depends on luminance, the intrinsic spatial frequency of the network becomes a function of luminance. Stimuli whose spatial frequency lies near this intrinsic frequency elicit the strongest responses, giving rise to a preferred spatial frequency that shifts with background luminance.

In this framework, changes in luminance do not simply rescale responses. Instead, they move the network between dynamical regimes by altering the balance of excitation and inhibition, which in turn shifts the intrinsic spatial frequency of the network and thus reorganizes spatial selectivity.

## Results

### Results of Modelling

We first examined how spatial-frequency selectivity arises from local interactions within the recurrent excitatory–inhibitory network (Fig. 1A–B). Varying the spatial wavenumber of the external stimulus (Fig. 1E) revealed that the response amplitude of the excitatory population depends strongly on stimulus wavenumber and reaches a maximum near the intrinsic spatial wavenumber of the circuit (Fig. 1H). This behavior is analogous to resonance phenomena in physics, in which responses reflect the interaction between external input and intrinsic dynamics. In such networks, local interactions give rise to a characteristic spatial coupling, illustrated by the impulse response of the circuit in Fig. 1C–D.

We define the spatial wavenumber at which the excitatory response coefficient *ξ(k, L*_*B*_) reaches its maximum as the resonance spatial wavenumber *k*_*r*_. Its corresponding ordinary spatial frequency is f_*r*_ = *k*_*r*_/(2*π*). We denote the experimentally measured preferred spatial frequency by f_pref_. The framework predicts that f_pref_ should track f_*r*_ because maximal perceptual sensitivity occurs when the stimulus spatial frequency approaches the resonance frequency of the recurrent circuit. The emergence of a well-defined resonance reflects the interaction between the spatial frequency of the stimulus and the intrinsic coupling structure of the recurrent circuit. Waves generated across the spatial extent of the stimulus superpose and produce constructive or destructive interference. These interactions selectively amplify the response to grating stimuli whose spatial frequency is close to the circuit’s intrinsic spatial frequency.

We next examined how background luminance modulates baseline activity in the circuit in the absence of spatial pattern. As shown in Fig. 1F, excitatory and inhibitory baseline activities vary non-linearly with luminance, with both responses increasing and saturating at higher luminance levels. Because the excitatory and inhibitory populations do not increase proportionally, the effective balance between excitation and inhibition shifts systematically with luminance, altering the operating state of the recurrent network. These changes define the operating point of the network (see Materials and Methods, “Effect of luminance”).

As shown in Fig. 1G, both the intrinsic and resonance spatial wavenumbers vary nonlinearly with mean luminance. At low luminance, both wavenumbers remain approximately constant. As luminance increases, the resonance wavenumber undergoes a sharp transition over a restricted luminance range and then approaches a higher plateau.

The transition is not determined by the absolute levels of excitation or inhibition alone. Rather, it emerges because changes in mean luminance alter the operating state of the recurrent network through changes in the effective excitatory–inhibitory balance. Stimulus frequencies near this intrinsic frequency elicit the strongest responses and therefore correspond to maximal sensitivity.

Together, these results suggest that the observed nonlinear dependence of preferred spatial frequency on luminance reflects changes in the operating state of the recurrent network, arising from luminance-dependent modulation of the effective excitatory–inhibitory balance.

### Results of psychophysical experiments

Contrast sensitivity functions were measured across a wide range of mean luminance *L*_*υ*_, revealing band-pass sensitivity to spatial frequency at each luminance level (Fig. 2D). Preferred spatial frequency f_pref_ was defined operationally as the spatial frequency at which the fitted phenomenological CSF 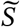 (f), defined by Eq. (21), reached its maximum (see Materials and Methods for the fitting procedure and Eq. (17)), yielding a consistent measure across luminance.

**Figure 2:**
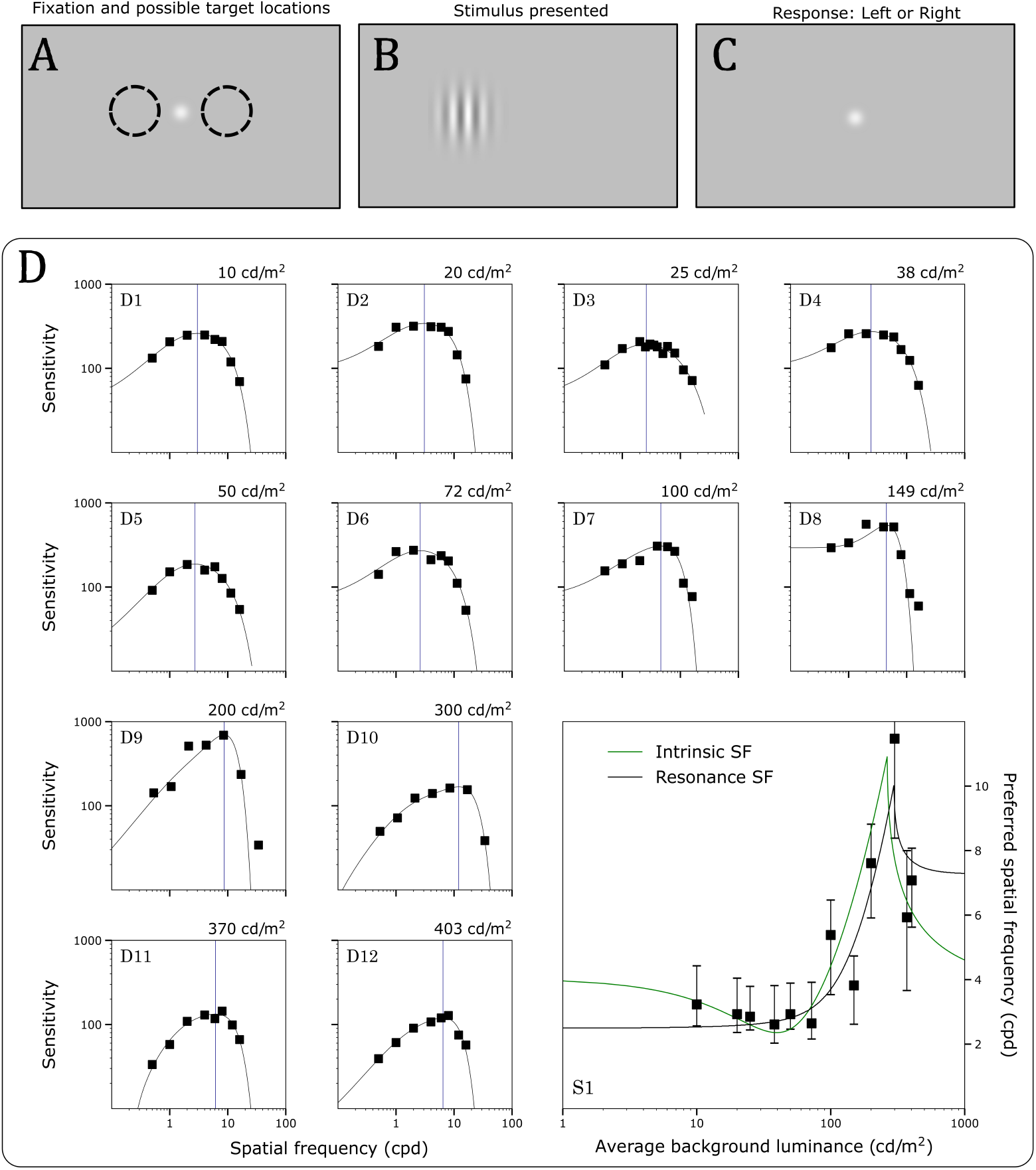
Trial structure and results in one subject. (**A–C**) Task and stimulus. A spatially periodic luminance grating was presented at one of two possible locations, to the left or right of a central fixation point (A). An example stimulus is shown in B. Subjects reported its location: left or right (C). (**D**) Contrast sensitivity functions measured at twelve levels of mean luminance *L*_*υ*_ in one subject. In each small panel, sensitivity is plotted as a function of spatial frequency. Solid curves are the fitted phenomenological CSF (Materials and Methods, Eq. (21)). The vertical lines mark the preferred spatial frequencies f_pref_, defined as the frequencies at which the fitted sensitivity functions are maximal (Eq. (17)), at the mean luminances displayed at the top right of each panel. In the bottom-right panel, preferred spatial frequencies are plotted as a function of mean luminance for the same subject. Black points show the estimated preferred spatial frequencies, and vertical bars indicate the 2.5th and 97.5th percentiles of the retained resampled preferred-spatial-frequency distribution. The black curve shows the fitted resonance spatial frequency f_*r*_(*L*_*B*_) of the recurrent-network model (Materials and Methods, Eq. (16)), and the green curve shows a separate fit of the model’s intrinsic spatial frequency f_*n*_(*L*_*B*_) (Materials and Methods, Eq. (20)). The two model quantities were fitted independently to the measured preferred spatial frequencies, as described in Materials and Methods.

In each subject, preferred spatial frequency varied systematically with luminance, remaining approximately constant at low luminance and shifting toward higher spatial frequencies over a restricted luminance range (Fig. 2D). This transition was reproducible across the four subjects tested (Fig. 3), indicating that it was not confined to a single observer. At higher luminance levels, preferred spatial frequency did not increase monotonically and in some subjects declined, suggesting that the relationship between luminance and preferred spatial frequency is not a simple monotonic scaling. Across subjects, the magnitude of the preferred spatial frequency shift spanned several cycles per degree and was confined to a narrow luminance interval, indicating a rapid reorganization rather than a gradual shift.

**Figure 3:**
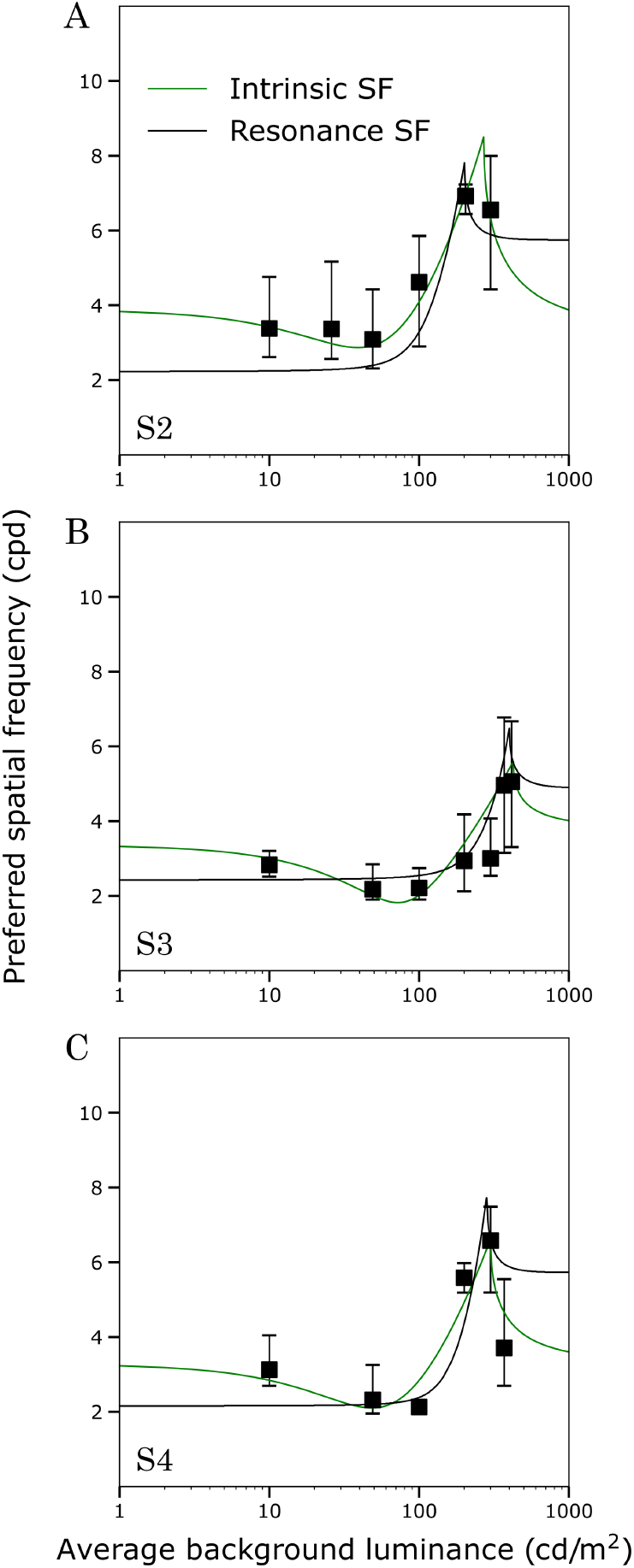
Preferred spatial frequency across luminance in three additional subjects. Preferred spatial frequencies f_pref_ are plotted as a function of mean luminance *L*_*υ*_ for Subjects 2–4 (**A–C**), following the format of Fig. 2D. In each panel, the black curve shows the model resonance spatial frequency f_r_(*L*_*B*_) and the green curve shows the separately evaluated intrinsic spatial frequency f_*n*_(*L*_*B*_). Black points show the estimated values of f_pref_, and vertical bars indicate the 2.5th and 97.5th percentiles of the retained resampled preferred-spatial-frequency distribution. The two model curves were evaluated separately and should not be interpreted as joint or unique parameter estimates.

Together, these measurements establish a robust luminance-dependent reorganization of spatial-frequency preference that the modelling framework seeks to explain. These measurements characterize the transition and motivate examination of whether it reflects changes in the operating state of the recurrent network, including shifts in the effective balance between excitation and inhibition.

## Discussion

Several influential accounts have described how contrast sensitivity changes with luminance, but they differ in emphasis and level of explanation. Classical psychophysical studies attributed luminance-dependent changes in contrast sensitivity primarily to detectability: at low luminance, noise and changes in effective spatial integration limit sensitivity to fine spatial detail, whereas increasing luminance improves the reliability with which spatial information can be encoded [Van Nes et al., 1967, Kelly, 1972, De Valois et al., 1974]. Subsequent physiological work explained these effects in terms of normalization and gain-control mechanisms, in which neural responses are regulated as luminance changes and overall activity is rescaled. Together, these accounts successfully describe many properties of luminance-dependent contrast sensitivity. However, they do not explain why the spatial frequency at which sensitivity is maximal reorganizes with luminance, nor how such reorganization emerges from cortical circuitry.

In the present measurements, preferred spatial frequency exhibited a characteristic S-shaped dependence on luminance, indicating a nonlinear reorganization of spatial selectivity rather than a simple monotonic rescaling of sensitivity. Previous psychophysical and physiological accounts can explain the general tendency of contrast sensitivity functions to shift toward higher spatial frequencies as luminance increases [De Valois et al., 1974, Kelly, 1977]. However, they do not by themselves predict the sharp transition between luminance regimes observed in the present data. To our knowledge, previous studies typically employed relatively coarse luminance sampling and focused on overall sensitivity or the location of the contrast-sensitivity peak [Kelly, 1972, Van Nes et al., 1967], rather than tracking preferred spatial frequency across luminance at sufficient resolution to reveal this transition. These observations motivate the need for a mechanistic framework linking luminance-dependent changes in cortical state to changes in spatial selectivity. In the present framework, changes in mean luminance (*L*_*B*_) alter excitatory and inhibitory baseline activity, thereby modifying effective recurrent coupling within the network. These changes shift the intrinsic spatial frequency of the circuit (f_*n*_), which in turn gives rise to corresponding changes in preferred spatial frequency.

While the present framework accounts for the sharp S-shaped dependence through luminance-dependent changes in the operating state of the recurrent network, the physiological mechanisms that generate these changes in effective excitatory–inhibitory balance remain incompletely understood. Previous physiological and theoretical studies have suggested that changes in stimulus contrast can often be approximated as a rescaling of effective input drive through normalization and gain-control mechanisms [Carandini and Heeger, 2012, Ohzawa et al., 1985], whereas changes in mean luminance may alter receptive-field organization or the effective response kernel underlying sensory selectivity [Cox et al., 1999, De Valois et al., 1982, Rahimi-Nasrabadi et al., 2021, Hildreth and Koch, 1987]. Consistent with these observations, physiological measurements summarized in Fig. 4 show luminance-dependent cortical inhibition [Tucker and Fitzpatrick, 2006], while excitatory and inhibitory responses exhibit different relative scaling with optogenetic drive [Adesnik, 2018]. Within the present framework, such changes may be interpreted as changes in effective excitatory–inhibitory balance, providing a plausible physiological basis for the observed reorganization of spatial selectivity.

**Figure 4:**
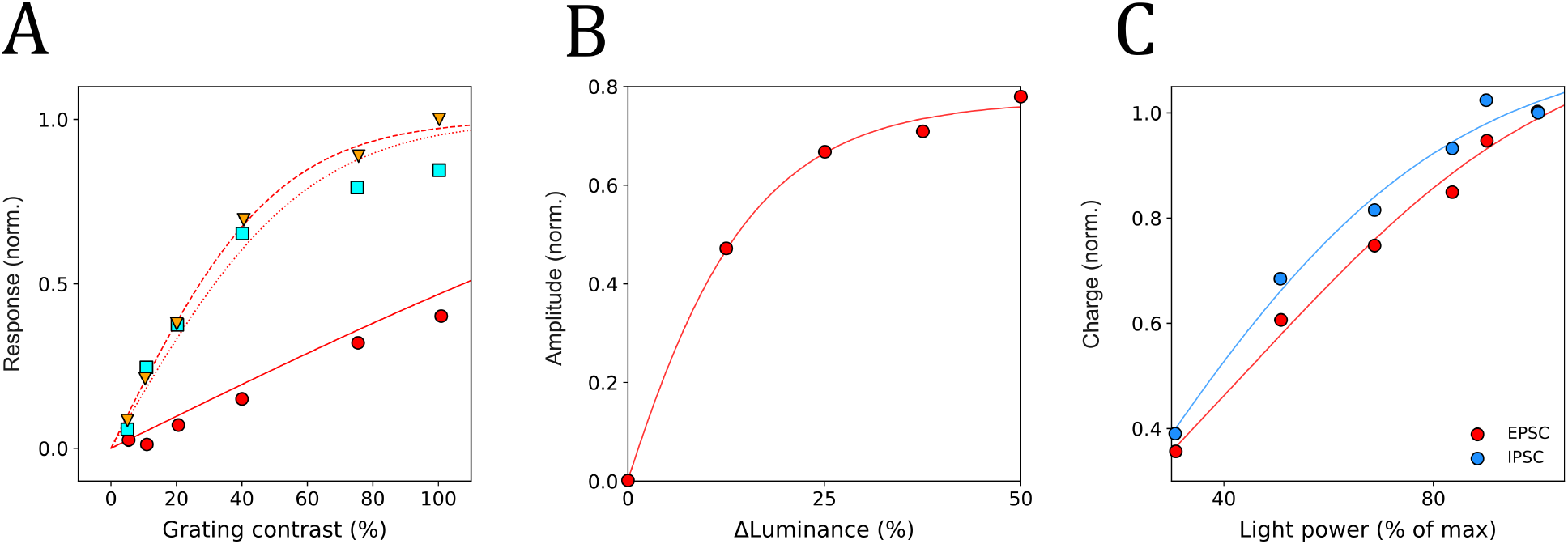
Comparison with previous neural response measurements. (**A**) Normalized responses of an LGN neuron to gratings of varying contrast at three stimulus diameters: 20° (red circles), 2° (cyan squares), and 0.5° (orange downward triangles), digitized from Fig. 2f of [Carandini and Heeger, 2012]; original data from [Bonin et al., 2005]. The corresponding recurrent excitatory–inhibitory model fits are shown as solid, dotted, and dashed red curves, respectively. (**B**) Population-averaged normalized cortical hyperpolarization as a function of luminance-step magnitude, replotted from [Tucker and Fitzpatrick, 2006]. Red circles show the published measurements and the solid red curve shows the model fit. (**C**) Population-averaged excitatory postsynaptic-current responses (EPSCs; red circles) and inhibitory postsynaptic-current responses (IPSCs; blue circles) in mouse primary visual cortex as a function of optogenetic stimulation power, replotted from [Adesnik, 2018]. The solid red and blue curves show the corresponding recurrent excitatory–inhibitory model fits. See Materials and Methods, “Curve fitting procedures,” for details of the fits.

As illustrated in Fig. 4A, the recurrent excitatory–inhibitory architecture used here reproduces classical normalization-like neural response profiles [Carandini and Heeger, 2012], suggesting that recurrent interactions implicated in normalization phenomena may also contribute to luminance-dependent changes in sensory tuning. Figure 4B further shows that larger luminance changes evoke progressively stronger cortical hyperpolarization [Tucker and Fitzpatrick, 2006], consistent with luminance-dependent recruitment of cortical inhibition. Complementing this observation, Fig. 4C demonstrates that excitatory and inhibitory responses both vary nonlinearly with optogenetic drive while differing in their relative scaling [Adesnik, 2018], showing that the effective balance between excitation and inhibition can vary with input drive. Together, the luminance-dependent inhibition in Fig. 4B and the drive-dependent excitatory–inhibitory scaling in Fig. 4C are consistent with luminance-dependent changes in the operating state of recurrent cortical circuits. Taken together, these physiological observations suggest that luminance changes the operating state of recurrent cortical circuits, providing a plausible framework through which luminance may reorganize spatial selectivity.

The present framework differs from conventional gain-control formulations in an important respect. Whereas gain-control accounts primarily describe how luminance modulates response amplitude, the present framework predicts that luminance can also alter the spatial frequency at which sensitivity is maximal. In the model, this occurs because luminance-dependent changes in excitatory–inhibitory balance shift the intrinsic spatial frequency of the recurrent network, producing corresponding changes in preferred spatial frequency. The observed reorganization of spatial selectivity therefore emerges as a consequence of changes in circuit state rather than from an explicit luminance-dependent tuning rule.

The proposed role of an intrinsic spatial frequency is consistent with growing evidence that cortical activity propagates through spatially distributed excitatory–inhibitory interactions rather than being determined solely by local receptive-field structure. Experimental and theoretical studies have demonstrated travelling and standing wave dynamics in visual cortex, suggesting that sensory processing is shaped by recurrent lateral interactions distributed across cortical space [Muller et al., 2018, Grabot et al., 2025, Sato et al., 2012, Bressloff, 2012, Gepshtein et al., 2022]. Within such frameworks, band-pass selectivity and spatial resonance emerge naturally from the coupling structure of cortical circuits [Daugman, 1985, Olshausen and Field, 1997, Bell and Sejnowski, 1997, Yamins et al., 2014]. From this perspective, luminance-dependent changes in excitatory–inhibitory balance may alter the intrinsic spatial scale of cortical activity, thereby shifting the spatial frequency at which sensitivity is maximal.

More broadly, these findings suggest that sensory selectivity may reflect the dynamical organization of recurrent cortical circuits as well as fixed feature-detector properties. Within the present framework, preferred spatial frequency is not externally imposed or entirely fixed, but emerges from the intrinsic spatial scale of recurrent interactions. Feature selectivity may therefore be state-dependent rather than static, with luminance altering not only response strength but also the operating state in which selectivity is expressed. Sensory tuning thus reflects an interaction between circuit architecture and the current dynamical state of the system. Stability and selectivity may consequently arise from the same recurrent interactions: the circuitry that stabilizes cortical activity also constrains the spatial tuning of the network.

In conclusion, the combined psychophysical and modelling results support a unified account in which luminance-dependent changes in preferred spatial frequency emerge from recurrent cortical dynamics. More generally, these findings suggest that sensory feature selectivity may emerge from the dynamical state of recurrent cortical circuits rather than solely from fixed receptive-field properties. This framework extends conventional gain-control descriptions of response amplitude by providing a mechanistic account of how luminance-dependent changes in cortical state can reorganize sensory selectivity. Whether similar state-dependent reorganizations extend to other sensory dimensions or adaptation paradigms remains an important question for future work.

## Materials and Methods

### Modelling methods

#### Model setup

In the simplified wave formalism presented below, neural activity in a one-dimensional homogeneous and isotropic chain of excitatory-inhibitory neurons is described by [Gepshtein et al., 2022]

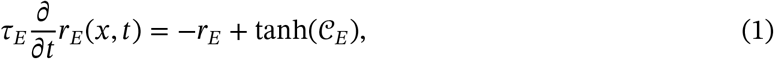

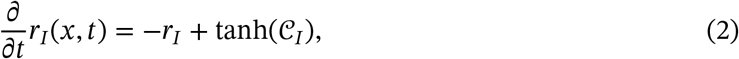

where the population inputs are given by

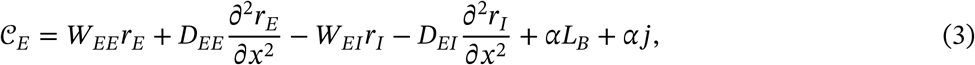

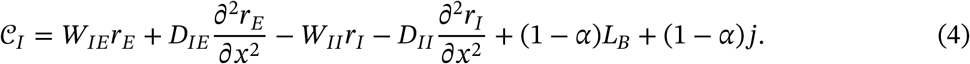

These equations describe the non-spontaneous activities *r*_*E*_(*x, t*) and *r*_*I*_(*x, t*) of the excitatory and inhibitory populations at cortical location *x* and time *t*. For the purposes of modelling visual processing, we assume that there is one-to-one spatial mapping between the stimulus coordinate *x* and the coordinate of the neural tissue. Under this mapping, a spatially extended stimulus can generate neural waves, whose interactions shape the population response across locations and over time.

The positive coupling parameters *W* and diffusion coefficients *D* describe the strength of interactions within and across locations in the excitatory and inhibitory populations (see Fig. 1(A)–(B)). The coupling weights w_*PQ*_and 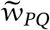 denote, respectively, within-motif and between-motif connections, where *P, Q ∈* {*E, I*}. With the unit motif spacing used here, the corresponding continuum parameters are

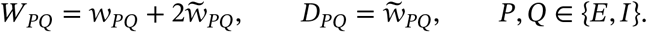

The coupling weights w_*PQ*_ and 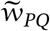 are defined in the prototype model, Eq. (1) of the Supplementary Materials in [Gepshtein et al., 2022].

The parameter *τ*_*E*_ denotes the excitatory time constant relative to the inhibitory time constant, which is scaled to unity.

Here, *j*(*x, t*) denotes the stimulus-driven external current, while *L*_*B*_ denotes the spatially and temporally constant background input characterising the mean luminance. The total external currents to the excitatory and inhibitory populations are therefore

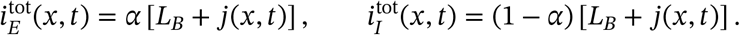

The parameter *α* controls how the total visual input is divided between the excitatory and inhibitory populations.

#### Effect of luminance

To examine how background luminance modifies the operating state of the network, we assume that the stimulus-driven current is small, |*j*(*x, t*)| ≪ 1, and apply a perturbation expansion that separates background activity from stimulus-driven activity:

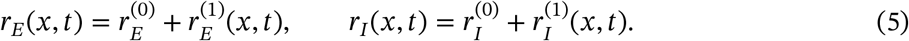

Substituting Eq. (5) into Eqs. (1)–(2) and collecting terms of equal order yields separate equations for the background activity and the stimulus-driven response. The zeroth-order terms 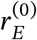and 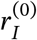 describe spatially uniform background activity and satisfy two algebraic transcendental equations

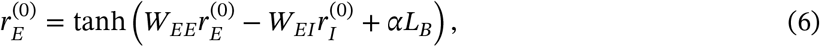

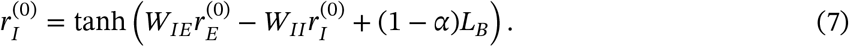

#### Linearization and renormalized parameters

Steady-state wave solutions appear in the first-order components 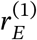 and 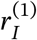 defined in Eq. (5). Because the spatially uniform background input *L*_*B*_ is incorporated into the zeroth-order activities 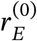and 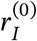, the first-order equations are driven only by the stimulus-dependent perturbations. These components satisfy the coupled linearized partial differential equations

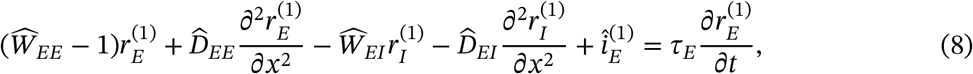

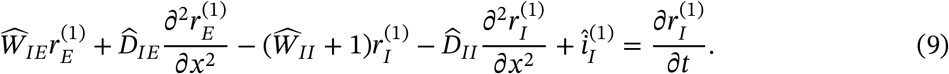

where the renormalized parameters incorporate the background activity through the luminance-dependent terms, 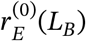 and 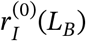 as:

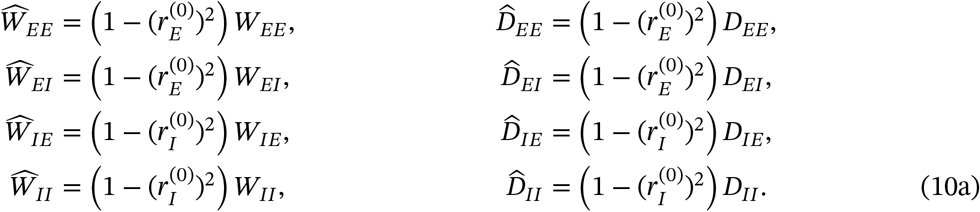

The first-order stimulus currents are 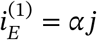 and 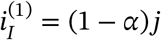. The same local gain factors that renormalize the interaction parameters also scale these first-order stimulus currents, giving

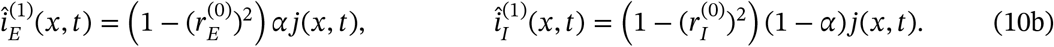

For each value of background luminance *L*_*B*_, Eqs. (6)–(7) were first solved numerically to determine the homogeneous steady-state activities 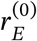 and 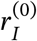. These values define the renormalized interaction parameters that characterize the operating state of the recurrent network. Two complementary linear-response problems were then considered using Eqs. (8)–(9) for spatially periodic forcing and localized point stimulation.

These complementary analyses characterize different properties of the network. The point-stimulation problem determines the intrinsic spatial wavenumber *k*_*n*_ from the Green’s-function response of the recurrent network. The periodically forced calculation determines the resonance spatial wavenumber *k*_*r*_ from the excitatory response coefficient *ξ(k, L*_*B*_).

#### Spatially periodic stimulus

To determine the resonance properties of the recurrent network under periodic forcing, the external input to Eqs. (1)–(2) was assumed to be spatially periodic,

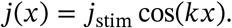

where *j*_stim_ is the stimulus amplitude and *k* is the stimulus spatial wavenumber. When *x* is expressed in degrees of visual angle, *k* has units of radians per degree. The corresponding ordinary spatial frequency, measured in cycles per degree, is

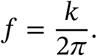

We denote the resonance and intrinsic spatial wavenumbers by *k*_*r*_ and *k*_*n*_, respectively. For comparison with the psychophysical measurements, these are expressed as ordinary spatial frequencies,

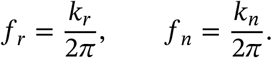

Because the stimulus is time independent, we henceforth write *j*(*x*) in place of *j*(*x, t*). For this spatially periodic input, the renormalized first-order inputs can be written as

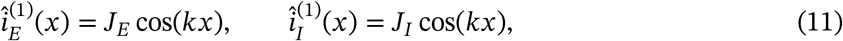

where the corresponding input amplitudes are

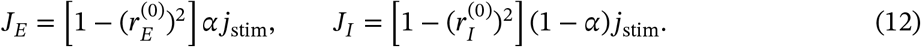

We seek steady-state solutions to Eqs. (8)–(9), and therefore adopt the notation 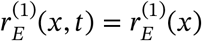 and 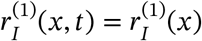, with

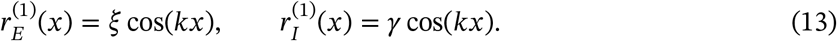

Substituting Eq. (13) and the corresponding spatially periodic input currents into Eqs. (8)–(9), and cancelling the common factor cos(*kx*), gives

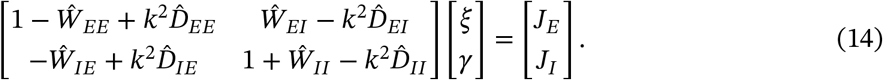

Solving Eq. (14) gives the excitatory response coefficient

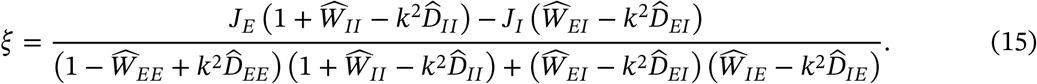

### Resonance spatial wavenumber

For each background luminance *L*_*B*_, the resonance spatial wavenumber *k*_*r*_ is defined as the value of *k* at which the excitatory response coefficient *ξ(k, L*_*B*_) attains its maximum over the wavenumber interval considered. At a smooth interior maximum,

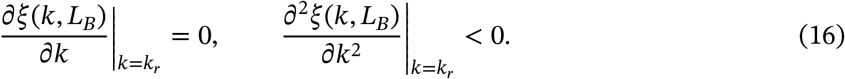

In the numerical implementation, trial ordinary spatial frequencies f were converted internally to wavenumbers using *k* = 2*π*f, and the excitatory response coefficient *ξ(2π*f, *L*_*B*_) was evaluated on the frequency grid used in the numerical continuation. The grid frequency at which *ξ was lar*gest was recorded as f_*r*_. No absolute value was applied to *ξ*.

#### *Pr*eferred spatial frequency

The experimentally measured preferred spatial frequency f_pref_ is defined from a smooth fitted representation of the contrast-sensitivity function, denoted by 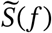. The preferred spatial frequency is the value of f at which 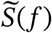 reaches its maximum and therefore satisfies

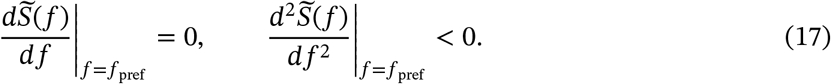

Thus, *k*_*r*_ is the model resonance spatial wavenumber, whereas f_pref_ is the experimentally measured preferred spatial frequency. Their comparison is made in ordinary spatial-frequency units through f_*r*_ = *k*_*r*_/(2*π*). Within the proposed framework, f_pref_ is expected to track f_*r*_.

#### Intrinsic spatial wavenumber

The intrinsic spatial wavenumber describes the natural spatial scale at which activity propagates within the recurrent network in the absence of external stimulation. Because it depends only on the renormalized interaction parameters, it is determined entirely by the operating state of the circuit.

Setting the external stimulus perturbation to zero reduces Eqs. (8)–(9) to the corresponding homogeneous system. To obtain the intrinsic spatial wavenumber, we seek solutions for *x* ≥ 0 of the form

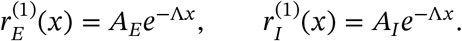

For the point-stimulation problem at *L*_*B*_ = 0, the damped spatial eigenvalue is written

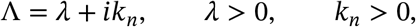

where *λ* is the spatial decay rate and *k*_*n*_ is the intrinsic spatial wavenumber of the oscillatory Green’s-function response.

For the luminance-dependent extension, the renormalized interaction parameters produce the two algebraic roots

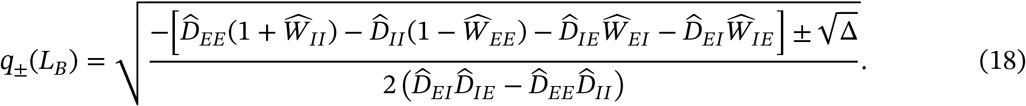

where the discriminant Δ is

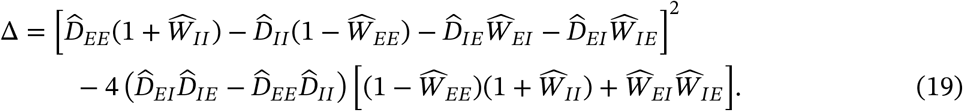

Here, *q*_±_ are complex spatial-wavenumber roots written in the *e*^*iqx*^ convention, rather than the decay eigenvalue Λ used in the Green’s-function representation *e*^−Λ*x*^. Following the continuation convention used in the model, the luminance-dependent intrinsic spatial wavenumbers are defined by

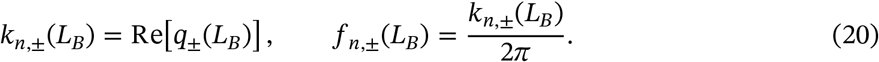

Under this convention, Re[*q*_±_] gives the oscillatory spatial wavenumber; the imaginary part describes spatial attenuation and is not used as the fitted spatial-frequency quantity.

Both real-part branches are shown in Fig. 1G. The quantitative intrinsic-frequency curves in Figs. 2–3 use the minus-sign branch,

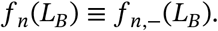

Only finite values of Re[*q*_±_(*L*_*B*_)] were used.

Equation (18) yields two algebraic branches, corresponding to the plus and minus signs before 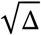. Thus, *k*_*r*_ and *k*_*n*_ denote model spatial wavenumbers, while f_*r*_ and f_*n*_ denote their corresponding ordinary spatial frequencies. The experimentally measured quantity is f_pref_. Comparisons with the psychophysical data are made using f_pref_, f_*r*_, and f_*n*_, all expressed in cycles per degree.

### Experimental methods

#### Participants

Four subjects (ages 20–25 years, all male) with normal or corrected-to-normal vision took part in the study. All participants received extensive training on the task prior to data collection and were compensated for their time. Experiments were conducted across three months. Each participant completed a series of contrast sensitivity function (CSF) measurements across a range of spatial frequencies (minimum of 7 spatial frequency conditions) and adaptation luminance levels (minimum of six, starting at 10 cd/m^2^). All procedures were approved by the Loughborough University Ethics Review Sub-Committee (Project ID 17492), and written informed consent was obtained from all participants.

#### Apparatus

All experiments were conducted in a dark room (<0.5 cd/m^2^ mean luminance) using a 27-inch 4K UHD monitor (3840 × 2160 resolution, 10-bit grayscale, 60 Hz refresh rate). Luminance calibration was performed using a Konica Minolta LS-150 luminance meter, ensuring accurate display output in cd/m^2^. Stimuli were generated and controlled via custom Python software (Pyglet / PsychoPy). During testing, the display area was limited to 2560 × 1440 pixels. Participants viewed stimuli binocularly from a distance of 100 cm, with head position stabilized by a chin-and-head rest.

#### Stimuli

The CSF stimulus was a Gaussian-windowed spatially periodic grating with a 1° spatial spread. For each spatial-frequency condition, multiple independent staircases were run. Gratings appeared randomly to the left or right of fixation (±0.5° displacement from fixation to grating center).

#### Procedure

Each trial began with a dim fixation dot with a Gaussian luminance profile, presented at the centre of the screen for 500 ms (Fig. 2A). The fixation dot was then removed and the grating was presented for 300 ms either to the left or right of the previous fixation location (Fig. 2B). Following stimulus offset, the fixation dot reappeared and remained visible until the participant responded. Participants used the left or right arrow key to report the perceived stimulus location (Fig. 2C). The response initiated the next trial.

Contrast detection thresholds were determined using a four-down, one-up adaptive staircase (step sizes: Δ^+^=0.36, Δ^−^=0.30294 log units). The procedure terminated after 10 reversals. For each spatial-frequency condition, up to four independent staircases were completed. Threshold estimation and the subsequent preferred-spatial-frequency analysis were performed using the resampling procedure described below. Preferred spatial frequency was determined as follows. Contrast-sensitivity estimates across spatial frequencies were fitted using a phenomenological exponential-plus-Gaussian function inspired by established analytical descriptions of the contrast sensitivity function [Watson and Ahumada, 2005, Campbell et al., 1966, Watson, 2006]:

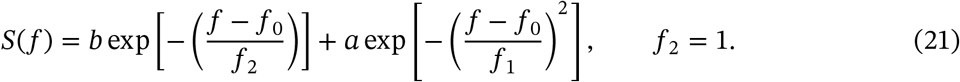

The parameter f_2_ was fixed at 1 cycle/degree and was therefore not optimized. The first exponential term is intentionally neither squared nor expressed using an absolute value; Eq. (21) reproduces the function implemented in the fitting code. The function was used solely as a phenomenological interpolant, and its parameters were not assigned physiological meaning.

Preferred spatial frequency was defined as the spatial frequency at which the fitted function attained its maximum. Although different parameter combinations may produce closely similar fitted contrast-sensitivity functions, the quantity of interest was the location of this maximum rather than the individual parameter values.

#### Resampling estimation of preferred spatial frequency

Each adaptive staircase was terminated after 10 reversals. To propagate uncertainty in the staircase threshold through the estimation of preferred spatial frequency, the reversal data were resampled. On each resampling iteration, eight of the 10 reversals obtained for each completed staircase were randomly selected without replacement and used to estimate its contrast threshold. Threshold estimates from the available staircases were then combined for each spatial-frequency condition and converted to contrast sensitivity, yielding one resampled contrast-sensitivity function (CSF).

For each subject and luminance condition, 10,000 resampling iterations were performed. On every iteration, the phenomenological CSF function (Eq. (21)) was fitted independently to the resampled contrast-sensitivity estimates, and the preferred spatial frequency f_pref_ was obtained as the spatial frequency at which the fitted function reached its maximum. This procedure therefore generated a distribution of 10,000 candidate preferred-spatial-frequency estimates for each subject and luminance condition.

Fit quality was assessed using the normalized root-mean-square error (NRMSE). An NRMSE threshold of 0.3 was adopted as an empirical quality-control criterion after visual inspection of representative fits indicated that values above this threshold generally corresponded to inadequate representations of the measured CSF. Fits satisfying 0 ≤ NRMSE ≤ 0.3 were therefore retained.

Inspection of the retained resampled distributions occasionally revealed a small, isolated group of preferred-spatial-frequency estimates at or below 1.05 cycles/degree, clearly separated from the dominant approximately bell-shaped distribution. These low-frequency estimates occurred when the maximum of the fitted CSF collapsed to the lower end of the evaluated frequency range and were therefore treated as fitting failures rather than reliable estimates of the CSF peak. They were excluded before calculation of the summary statistics. The reported preferred spatial frequency was the mean of the remaining f_pref_ distribution, and the error bars were defined by its 2.5th and 97.5th percentiles, corresponding to a 95% percentile interval.

### Curve fitting procedures

Several figures include fitted curves. The fitting procedures differed according to the purpose of each analysis and are summarized below. Unless otherwise stated, nonlinear least-squares optimization was performed using the Trust Region Reflective (TRF) algorithm implemented in Python’s `scipy.optimize.curve_fit`. All fitted curves are intended to summarize the data or evaluate model consistency and should not be interpreted as unique parameter estimates.

#### Empirical contrast sensitivity functions (Figure 2D)

For each measured contrast sensitivity function (CSF), an empirical function was fitted (Eq. (21)) to the psychophysical data using nonlinear least-squares optimization (Python `scipy.optimize.curve_fit`). No constraints were imposed on the parameter values during optimization, allowing the fitting routine to determine the parameter combination that minimized the residual sum of squares.

The fitted function served only as a smooth interpolation of the measured CSF and was not interpreted mechanistically. Consequently, the fitted parameters themselves were not assigned physiological meaning. Instead, the preferred spatial frequency (PSF) was defined as the spatial frequency corresponding to the maximum of the fitted CSF. All subsequent analyses were performed using this estimated peak location rather than the individual fitting parameters.

An example fit for Subject 1 is shown in Fig. 2D. The fitted parameter values are omitted because the empirical CSF function was used solely as a smooth interpolation of the measured data, and no physiological interpretation is attached to the individual parameters.

#### Model fits to psychophysical measurements (Figures 2–3)

The experimentally measured preferred spatial frequencies were compared with two distinct quantities derived from the recurrent network model: the intrinsic spatial frequency f_*n*_ = *k*_*n*,−_/(2*π*) and the resonance spatial frequency f_*r*_ = *k*_*r*_ / (2*π*). The two quantities were fitted separately to the psychophysical measurements.

Experimental mean luminance, denoted *L*_*υ*_, was supplied to the fitting procedure in physical units of cd/m^2^. No preliminary linear conversion to arbitrary model units was applied. Instead, each measured luminance value was mapped directly to the model input according to

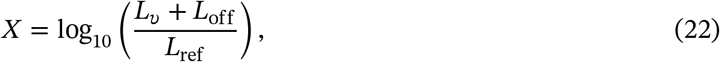

where *L*_off_ = 100 cd/m^2^ and *L*_ref_ = 1 cd/m^2^, and *L*_*υ*_ denotes experimental luminance in cd/m^2^. The reference luminance renders the argument of the logarithm dimensionless and is implicit in the numerical implementation. The offset and the multiplicative scale factor, which was fixed at 1, were not optimized during fitting.

The transformed luminance input was divided between the excitatory and inhibitory populations according to

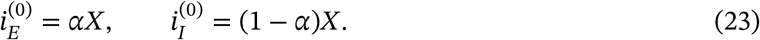

The model curves in Figs. 2 and 3 were therefore evaluated by applying this transformation directly to the measured luminances. The logarithmic horizontal axes in these figures display the original luminance values in cd/m^2^ and represent only a plotting convention, not an additional transformation of the model input.

For the intrinsic spatial-frequency fit, the transformed input *X* entered the analytical expression for *k*_*n*_ through the excitatory and inhibitory background-input terms. The recurrent and lateral coupling strengths, the excitatory input fraction *α*, and the initial background-activity parameters were optimized by nonlinear least squares. The resulting parameter set was then used to evaluate f_*n*_(*L*_*B*_) across the measured luminance range.

The minus-sign root was evaluated as *k*_*n*,−_(*L*_*B*_) = Re[*q*_−_(*L*_*B*_)] and converted to ordinary spatial frequency using f_*n*_(*L*_*B*_) = *k*_*n*,−_(*L*_*B*_)/(2*π*).

For each luminance level, trial frequencies f were converted to wavenumbers using *k* = 2*π*f. The excitatory response coefficient *ξ(2π*f, *L*_*B*_) was evaluated over the continuation grid, and the frequency giving the largest value of *ξ was rec*orded as f_*r*_(*L*_*B*_). No absolute value was applied.

The f_*n*_ and f_*r*_ curves were obtained from separate parameter optimizations. They therefore represent separate evaluations of whether the intrinsic and periodically forced responses of the recurrent network can reproduce the measured luminance dependence of preferred spatial frequency.

#### Model fits to published physiological data (Fig. 4)

The recurrent excitatory–inhibitory framework was also fitted to previously published physiological measurements shown in Fig. 4.

For Fig. 4A, seven of the eight coupling weights were held fixed, while a single coupling weight was varied to reproduce the different experimental stimulus conditions. This procedure tested whether variation of a single network interaction could account for the family of normalization-like response curves observed experimentally.

For Fig. 4B, the model was fitted separately to the population-averaged cortical hyperpolarization measurements. The recurrent excitatory–inhibitory model was numerically integrated to steady state for each luminance-step condition, after which scaling and offset parameters were used to map the model response onto the normalized experimental measurements.

For Fig. 4C, the excitatory and inhibitory responses were fitted simultaneously to the published EPSC and IPSC measurements. The recurrent excitatory–inhibitory model was numerically integrated to steady state for each optogenetic-stimulation-power condition. Shared recurrent coupling parameters were estimated by minimizing the combined residual error across the excitatory and inhibitory datasets, with linear scaling and offset parameters used to map the model responses onto the normalized experimental measurements.

