## supplementary materials for "Intrinsic Organization of Contrast Sensitivity in Human Vision"

This Supporting Information expands the mathematical derivations and fitting procedures used in the main manuscript. It is intended to make explicit the connection between the discrete recurrent excitatory–inhibitory (E–I) network, its continuum representation, the luminance-dependent linearization, the periodically forced response, and the intrinsic spatial modes of the network. The notation follows the current main manuscript. The supplementary equations are self-contained and are numbered independently as Eqs. (S1), (S2), ....

#### S1 Mathematical framework and notation

##### S1.1 Principal quantities

The model describes the activities of interacting excitatory and inhibitory populations distributed along a one-dimensional cortical coordinate $x$. The activity variables $r_E(x,t)$ and $r_I(x,t)$ are dimensionless population-level activities. The inhibitory time constant is used as the reference time scale, and the excitatory time constant relative to this reference is denoted by $\tau_E$.

For clarity, the main symbols used below are summarized in Table S1.

##### S1.2 From the discrete E–I chain to the continuum equations

The starting point is a chain of identical E–I motifs coupled to their nearest neighbours. At motif position $l$, the excitatory and inhibitory input fields can be written as

$$\begin{aligned}\mathcal{C}_E(l) = w_{EE} r_E(l) + \widetilde{w}_{EE}[r_E(l+1) + r_E(l-1)] \\ - w_{EI} r_I(l) - \widetilde{w}_{EI}[r_I(l+1) + r_I(l-1)] + i_E^{\text{tot}}(l,t),\end{aligned} \tag{S1}$$

$$\begin{aligned}\mathcal{C}_I(l) = w_{IE} r_E(l) + \widetilde{w}_{IE}[r_E(l+1) + r_E(l-1)] \\ - w_{II} r_I(l) - \widetilde{w}_{II}[r_I(l+1) + r_I(l-1)] + i_I^{\text{tot}}(l,t).\end{aligned} \tag{S2}$$

Table S1: Principal notation used in the supplementary derivations.

| Symbol | Meaning |
|---|---|
| $x, t$ | Cortical position and time. |
| $r_E, r_I$ | Excitatory and inhibitory population activities. |
| $w_{PQ}$ | Within-motif coupling weight from population $Q$ to population $P$, with $P, Q \in \{E, I\}$. |
| $\widetilde{w}_{PQ}$ | Between-motif (nearest-neighbour) coupling weight from population $Q$ to population $P$. |
| $W_{PQ}$ | Combined coupling parameter in the continuum representation. |
| $D_{PQ}$ | Spatial coupling (diffusion) coefficient in the continuum representation. |
| $L_B$ | Spatially and temporally uniform background input in the mathematical model, characterising the mean luminance. |
| $L_v$ | Experimentally measured mean luminance, in cd/m$^2$. |
| $j(x, t)$ | Stimulus-driven external input. |
| $i_P^{\text{tot}}$ | Total external input to population $P$, containing the background and stimulus components. |
| $\alpha$ | Fraction of the total visual input delivered to the excitatory population; $1-\alpha$ is delivered to the inhibitory population. |
| $r_P^{(0)}$ | Homogeneous background activity of population $P$. |
| $r_P^{(1)}$ | First-order stimulus-driven activity of population $P$. |
| $s_P$ | Local linear gain $1-(r_P^{(0)})^2$ about the background state. |
| $\widehat{W}_{PQ}, \widehat{D}_{PQ}$ | Luminance-dependent renormalized interaction parameters. |
| $\hat{i}_P^{(1)}$ | Gain-scaled first-order stimulus input. |
| $J_E, J_I$ | Harmonic amplitudes of the renormalized first-order excitatory and inhibitory inputs under spatially periodic forcing. |
| $k$ | Spatial wavenumber of an imposed periodic stimulus; $f = k/(2\pi)$ is the corresponding ordinary spatial frequency. |
| $k_r, f_r$ | Resonance spatial wavenumber and its ordinary spatial-frequency representation, $f_r = k_r/(2\pi)$. |
| $k_n, f_n$ | Intrinsic spatial wavenumber and its ordinary spatial-frequency representation, $f_n = k_n/(2\pi)$. |
| $f_{\text{pref}}$ | Experimentally measured preferred spatial frequency, in cycles per degree. |
| $\lambda$ | Spatial decay rate of the Green's-function response. |
| $\ell_n$ | Spatial wavelength associated with the intrinsic oscillatory response in Fig. 1D of the main text. |

Let $x = l\Delta x$ denote the continuum coordinate. For activity that varies smoothly from one motif to the next,

$$r_P(x \pm \Delta x) = r_P(x) \pm \Delta x \frac{\partial r_P}{\partial x} + \frac{(\Delta x)^2}{2} \frac{\partial^2 r_P}{\partial x^2} + \mathcal{O}((\Delta x)^3), \qquad P \in \{E, I\}. \tag{S3}$$

Adding the two neighbouring terms cancels the first spatial derivative,

$$r_P(x + \Delta x) + r_P(x - \Delta x) = 2r_P(x) + (\Delta x)^2 \frac{\partial^2 r_P}{\partial x^2} + \mathcal{O}((\Delta x)^4). \tag{S4}$$

Consequently, for each ordered population pair $PQ$,

$$W_{PQ} = w_{PQ} + 2\widetilde{w}_{PQ}, \qquad D_{PQ} = \widetilde{w}_{PQ}(\Delta x)^2, \qquad P, Q \in \{E, I\}. \tag{S5}$$

The model coordinate used in the main manuscript measures distance in units of the motif spacing, so that $\Delta x = 1$ and

$$D_{PQ} = \widetilde{w}_{PQ}. \tag{S6}$$

If $x$ is instead expressed explicitly in physical units, such as degrees of visual angle, the factor $(\Delta x)^2$ is retained in $D_{PQ}$. This continuum approximation yields

$$\mathcal{C}_E = W_{EE} r_E + D_{EE} \frac{\partial^2 r_E}{\partial x^2} - W_{EI} r_I - D_{EI} \frac{\partial^2 r_I}{\partial x^2} + i_E^{\text{tot}}(x,t), \tag{S7}$$

$$\mathcal{C}_I = W_{IE} r_E + D_{IE} \frac{\partial^2 r_E}{\partial x^2} - W_{II} r_I - D_{II} \frac{\partial^2 r_I}{\partial x^2} + i_I^{\text{tot}}(x,t). \tag{S8}$$

The total external inputs are partitioned between the two populations as

$$i_E^{\text{tot}}(x,t) = \alpha \left[L_B + j(x,t)\right], \qquad i_I^{\text{tot}}(x,t) = (1-\alpha)\left[L_B + j(x,t)\right]. \tag{S9}$$

The continuum Wilson–Cowan equations used in the main manuscript are then

$$\tau_E \frac{\partial r_E}{\partial t} = -r_E + \tanh(\mathcal{C}_E), \tag{S10}$$

$$\frac{\partial r_I}{\partial t} = -r_I + \tanh(\mathcal{C}_I). \tag{S11}$$

The inhibitory time constant has been scaled to unity, so $\tau_E$ is the excitatory time constant relative to the inhibitory population. This construction follows the spatially distributed Wilson–Cowan framework developed previously [Wilson and Cowan, 1972, 1973, Gepshtein et al., 2022].

### S2 Background state and luminance-dependent linearization

#### S2.1 Separation of background and stimulus-driven activity

The background input is spatially homogeneous, whereas the stimulus-driven input can vary in space and time. For $|j(x,t)| \ll 1$, we separate the population activity into a homogeneous background state and a small stimulus-driven perturbation,

$$r_E(x,t) = r_E^{(0)} + r_E^{(1)}(x,t), \qquad r_I(x,t) = r_I^{(0)} + r_I^{(1)}(x,t). \tag{S12}$$

For the derivation, it is useful to introduce a formal bookkeeping parameter $\varepsilon$ and write $j = \varepsilon j_1$ and $r_P = r_P^{(0)} + \varepsilon r_P^{(1)}$. The physical first-order equations are recovered by setting $\varepsilon = 1$ after collecting equal powers of $\varepsilon$.

At zeroth order, all spatial derivatives vanish and the background state satisfies

$$r_E^{(0)} = \tanh\left[W_{EE} r_E^{(0)} - W_{EI} r_I^{(0)} + \alpha L_B\right], \tag{S13}$$

$$r_I^{(0)} = \tanh\left[W_{IE} r_E^{(0)} - W_{II} r_I^{(0)} + (1-\alpha) L_B\right]. \tag{S14}$$

For each value of the background drive, these two coupled transcendental equations are solved numerically to determine the operating point $\left(r_E^{(0)}, r_I^{(0)}\right)$.

#### S2.2 Origin of the renormalized parameters

Let

$$\mathcal{C}_P = \mathcal{C}_P^{(0)} + \varepsilon\,\delta\mathcal{C}_P, \qquad P \in \{E, I\}. \tag{S15}$$

The first-order Taylor expansion of the nonlinear response function is

$$\tanh\Big(\mathcal{C}_P^{(0)} + \varepsilon\,\delta\mathcal{C}_P\Big) = \tanh(\mathcal{C}_P^{(0)}) + \varepsilon\,\mathrm{sech}^2(\mathcal{C}_P^{(0)})\delta\mathcal{C}_P + \mathcal{O}(\varepsilon^2). \tag{S16}$$

Because $r_P^{(0)} = \tanh(\mathcal{C}_P^{(0)})$, the local gain factor can be written as

$$s_E = \mathrm{sech}^2(\mathcal{C}_E^{(0)}) = 1 - \left(r_E^{(0)}\right)^2, \qquad s_I = \mathrm{sech}^2(\mathcal{C}_I^{(0)}) = 1 - \left(r_I^{(0)}\right)^2. \tag{S17}$$

Before introducing the hatted notation, the first-order equations are therefore

$$\tau_E \frac{\partial r_E^{(1)}}{\partial t} = -r_E^{(1)} + s_E\left[W_{EE} r_E^{(1)} + D_{EE}\frac{\partial^2 r_E^{(1)}}{\partial x^2} - W_{EI} r_I^{(1)} - D_{EI}\frac{\partial^2 r_I^{(1)}}{\partial x^2} + \alpha j\right], \tag{S18}$$

$$\frac{\partial r_I^{(1)}}{\partial t} = -r_I^{(1)} + s_I\left[W_{IE} r_E^{(1)} + D_{IE}\frac{\partial^2 r_E^{(1)}}{\partial x^2} - W_{II} r_I^{(1)} - D_{II}\frac{\partial^2 r_I^{(1)}}{\partial x^2} + (1-\alpha) j\right]. \tag{S19}$$

Thus, the luminance dependence of the linearized network enters through the local gains $s_E$ and $s_I$. We define

$$\begin{aligned} \widehat{W}_{EE} &= s_E W_{EE}, & \widehat{D}_{EE} &= s_E D_{EE}, & \widehat{W}_{EI} &= s_E W_{EI}, & \widehat{D}_{EI} &= s_E D_{EI}, \\ \widehat{W}_{IE} &= s_I W_{IE}, & \widehat{D}_{IE} &= s_I D_{IE}, & \widehat{W}_{II} &= s_I W_{II}, & \widehat{D}_{II} &= s_I D_{II}, \end{aligned} \tag{S20}$$

and the first-order stimulus inputs

$$i_E^{(1)} = \alpha j, \qquad i_I^{(1)} = (1-\alpha) j. \tag{S21}$$

The same gain factors scale these stimulus inputs,

$$\hat{i}_E^{(1)} = s_E i_E^{(1)} = s_E \alpha j, \qquad \hat{i}_I^{(1)} = s_I i_I^{(1)} = s_I (1-\alpha) j. \tag{S22}$$

With these definitions, the first-order dynamics become

$$(\widehat{W}_{EE} - 1) r_E^{(1)} + \widehat{D}_{EE}\frac{\partial^2 r_E^{(1)}}{\partial x^2} - \widehat{W}_{EI} r_I^{(1)} - \widehat{D}_{EI}\frac{\partial^2 r_I^{(1)}}{\partial x^2} + \hat{i}_E^{(1)} = \tau_E \frac{\partial r_E^{(1)}}{\partial t}, \tag{S23}$$

$$\widehat{W}_{IE} r_E^{(1)} + \widehat{D}_{IE}\frac{\partial^2 r_E^{(1)}}{\partial x^2} - (\widehat{W}_{II} + 1) r_I^{(1)} - \widehat{D}_{II}\frac{\partial^2 r_I^{(1)}}{\partial x^2} + \hat{i}_I^{(1)} = \frac{\partial r_I^{(1)}}{\partial t}. \tag{S24}$$

Thus, $\hat{i}_P^{(1)}$ denotes the first-order stimulus input after multiplication by the local slope of the nonlinear response function about the luminance-dependent background state; it does not represent an additional external input. The same gain-scaling interpretation applies to the hatted coupling and diffusion parameters.

### S3 Periodic forcing and resonance spatial wavenumber

#### S3.1 Steady response to a spatially periodic stimulus

For the periodic forcing calculation, the external input is taken to be spatially periodic,

$$j(x) = j_{\mathrm{stim}} \cos(kx), \tag{S25}$$

where $j_{\mathrm{stim}}$ is the stimulus amplitude and $k$ is the stimulus spatial wavenumber. When $x$ is expressed in degrees of visual angle, $k$ has units of radians per degree. The corresponding ordinary spatial frequency, measured in cycles per degree, is

$$f = \frac{k}{2\pi}. \tag{S26}$$

We denote the resonance and intrinsic spatial wavenumbers by $k_r$ and $k_n$, respectively, and their ordinary spatial-frequency representations by

$$f_r = \frac{k_r}{2\pi}, \qquad f_n = \frac{k_n}{2\pi}. \tag{S27}$$

Because the input is time independent, we henceforth write $j(x)$ in place of $j(x,t)$.

We seek steady-state first-order responses of the form

$$r_E^{(1)}(x) = \xi \cos(kx), \qquad r_I^{(1)}(x) = \gamma \cos(kx). \tag{S28}$$

Hence,

$$\frac{\partial^2}{\partial x^2} \cos(kx) = -k^2 \cos(kx). \tag{S29}$$

For this spatially periodic input, the renormalized first-order inputs can be written as

$$\hat{i}_E^{(1)}(x) = J_E \cos(kx), \qquad \hat{i}_I^{(1)}(x) = J_I \cos(kx), \tag{S30}$$

where the corresponding input amplitudes are

$$J_E = s_E \alpha j_{\mathrm{stim}}, \qquad J_I = s_I (1-\alpha) j_{\mathrm{stim}}. \tag{S31}$$

Equivalently, using Eq. (S17),

$$J_E = \left[1 - \left(r_E^{(0)}\right)^2\right] \alpha j_{\mathrm{stim}}, \qquad J_I = \left[1 - \left(r_I^{(0)}\right)^2\right] (1-\alpha) j_{\mathrm{stim}}. \tag{S32}$$

Substituting Eq. (S26) and the corresponding spatially periodic inputs into Eqs. (S23)–(S24), and cancelling the common factor $\cos(kx)$, gives

$$\mathbf{M}(k, L_B) \begin{bmatrix} \xi \\ \gamma \end{bmatrix} = \begin{bmatrix} J_E \\ J_I \end{bmatrix}, \tag{S33}$$

where

$$\mathbf{M}(k, L_B) = \begin{bmatrix} 1 - \widehat{W}_{EE} + k^2 \widehat{D}_{EE} & \widehat{W}_{EI} - k^2 \widehat{D}_{EI} \\ -\widehat{W}_{IE} + k^2 \widehat{D}_{IE} & 1 + \widehat{W}_{II} - k^2 \widehat{D}_{II} \end{bmatrix}. \tag{S34}$$

The hatted interaction parameters depend on $L_B$ through $r_E^{(0)}(L_B)$ and $r_I^{(0)}(L_B)$, and the amplitudes $J_E$ and $J_I$ inherit the same luminance-dependent gain factors.

Define

$$M_{11} = 1 - \widehat{W}_{EE} + k^2\widehat{D}_{EE}, \qquad M_{12} = \widehat{W}_{EI} - k^2\widehat{D}_{EI}, \tag{S35}$$

$$M_{21} = -\widehat{W}_{IE} + k^2\widehat{D}_{IE}, \qquad M_{22} = 1 + \widehat{W}_{II} - k^2\widehat{D}_{II}. \tag{S36}$$

The determinant is

$$\mathcal{D}(k, L_B) = M_{11}M_{22} - M_{12}M_{21}, \tag{S37}$$

or, equivalently,

$$\begin{aligned}\mathcal{D}(k, L_B) = {} & \left(1 - \widehat{W}_{EE} + k^2\widehat{D}_{EE}\right)\left(1 + \widehat{W}_{II} - k^2\widehat{D}_{II}\right) \\ & + \left(\widehat{W}_{EI} - k^2\widehat{D}_{EI}\right)\left(\widehat{W}_{IE} - k^2\widehat{D}_{IE}\right).\end{aligned} \tag{S38}$$

Solving the $2 \times 2$ system gives

$$\xi(k, L_B) = \frac{J_E M_{22} - J_I M_{12}}{\mathcal{D}(k, L_B)}, \tag{S39}$$

and

$$\gamma(k, L_B) = \frac{M_{11}J_I - M_{21}J_E}{\mathcal{D}(k, L_B)}. \tag{S40}$$

In expanded form,

$$\xi = \frac{J_E\left(1 + \widehat{W}_{II} - k^2\widehat{D}_{II}\right) - J_I\left(\widehat{W}_{EI} - k^2\widehat{D}_{EI}\right)}{\left(1 - \widehat{W}_{EE} + k^2\widehat{D}_{EE}\right)\left(1 + \widehat{W}_{II} - k^2\widehat{D}_{II}\right) + \left(\widehat{W}_{EI} - k^2\widehat{D}_{EI}\right)\left(\widehat{W}_{IE} - k^2\widehat{D}_{IE}\right)}. \tag{S41}$$

#### S3.2 Definition and numerical evaluation of resonance

For each fixed background state, the resonance spatial wavenumber $k_r$ is defined as the stimulus spatial wavenumber at which the signed excitatory response coefficient $\xi(k, L_B)$ reaches its maximum over the wavenumber domain considered. No absolute value is applied to $\xi$ in this definition. This is important because replacing $\xi$ by $|\xi|$ would alter the modelled response by converting large negative responses into positive response magnitudes.

At a smooth interior maximum,

$$\left.\frac{\partial \xi(k, L_B)}{\partial k}\right|_{k=k_r} = 0, \qquad \left.\frac{\partial^2 \xi(k, L_B)}{\partial k^2}\right|_{k=k_r} < 0. \tag{S42}$$

The derivative conditions characterize an interior local maximum. In the numerical implementation, trial ordinary spatial frequencies $f$ are converted to wavenumbers using $k = 2\pi f$, and $\xi(2\pi f, L_B)$ is evaluated on the frequency grid used for numerical continuation. The grid frequency at which $\xi$ is largest is recorded as $f_r$; equivalently, $k_r = 2\pi f_r$.

### S4 Intrinsic spatial organization of the recurrent network

The intrinsic spatial wavenumber is a property of the recurrent operator rather than of a particular periodic stimulus. Two complementary representations are useful. The Green's-function representation identifies the damped oscillatory spatial eigenvalue following localized stimulation, whereas the luminance-dependent calculation used for fitting obtains $k_n$ from the characteristic spatial-mode equation expressed in terms of the renormalized parameters.

#### S4.1 Point stimulation and the Green's-function representation

For a localized point input, the Green's function is the spatial impulse-response kernel of the linearized network. For population $P \in \{E, I\}$,

$$r_P^{(1)}(x) = \int_{-\infty}^{+\infty} j(x')\, G_P(x - x')\, dx'. \tag{S43}$$

For a point stimulus $j(x) = \delta(x)$, the response is proportional to the Green's function itself.

Away from the point of stimulation, the external perturbation vanishes and Eqs. (S23)–(S24) reduce, at steady state, to a homogeneous spatial system. Seeking solutions for $x \geq 0$ of the form

$$r_E^{(1)}(x) = A_E e^{-\Lambda x}, \qquad r_I^{(1)}(x) = A_I e^{-\Lambda x}, \tag{S44}$$

gives

$$\mathbf{M}_\Lambda \begin{bmatrix} A_E \\ A_I \end{bmatrix} = \begin{bmatrix} 0 \\ 0 \end{bmatrix}, \tag{S45}$$

with

$$\mathbf{M}_\Lambda = \begin{bmatrix} 1 - \widehat{W}_{EE} - \Lambda^2 \widehat{D}_{EE} & \widehat{W}_{EI} + \Lambda^2 \widehat{D}_{EI} \\ -\widehat{W}_{IE} - \Lambda^2 \widehat{D}_{IE} & 1 + \widehat{W}_{II} + \Lambda^2 \widehat{D}_{II} \end{bmatrix}. \tag{S46}$$

Non-trivial solutions require

$$\det \mathbf{M}_\Lambda = 0. \tag{S47}$$

For the damped oscillatory regime used in the Green's-function interpretation, the relevant spatial root is written

$$\Lambda = \lambda + i k_n, \qquad \lambda > 0, \qquad k_n > 0. \tag{S48}$$

The real part $\lambda$ sets the spatial decay rate and the imaginary component $k_n$ sets the intrinsic oscillatory spatial scale. The corresponding Green's functions have the generic form

$$G_P(x) = e^{-\lambda |x|} \left[ \Gamma_P \cos(k_n x) - \Delta_P \,\mathrm{sign}(x) \sin(k_n x) \right], \qquad P \in \{E, I\}, \tag{S49}$$

where $\Gamma_P$ and $\Delta_P$ are determined by the coupling parameters, the diffusion coefficients, and the matching conditions at the point source [Gepshtein et al., 2022].

The distinction between spatial wavelength and decay is explicit in this representation. In the notation of Fig. 1D of the main text, $\ell_n$ denotes the wavelength of the intrinsic oscillatory response, so that

$$k_n = \frac{2\pi}{\ell_n}, \qquad f_n = \frac{k_n}{2\pi} = \frac{1}{\ell_n}. \tag{S50}$$

The decay rate $\lambda$ independently determines how rapidly the response envelope decreases with distance.

#### S4.2 Luminance-dependent intrinsic modes

For the luminance-dependent analysis, $k_n$ is obtained from the homogeneous spatial-mode condition expressed in terms of the renormalized parameters. After setting the first-order external inputs to zero, we seek modes of the form

$$r_E^{(1)}(x) = A_E e^{iqx}, \qquad r_I^{(1)}(x) = A_I e^{iqx}, \tag{S51}$$

where $q$ may be complex. Non-trivial solutions require

$$\det \mathbf{M}(q, L_B) = 0. \tag{S52}$$

Writing $\mu = q^2$ gives the quadratic

$$a\mu^2 + b\mu + c = 0, \tag{S53}$$

where

$$a = \widehat{D}_{EI}\widehat{D}_{IE} - \widehat{D}_{EE}\widehat{D}_{II}, \tag{S54}$$

$$b = \widehat{D}_{EE}(1 + \widehat{W}_{II}) - \widehat{D}_{II}(1 - \widehat{W}_{EE}) - \widehat{D}_{IE}\widehat{W}_{EI} - \widehat{D}_{EI}\widehat{W}_{IE}, \tag{S55}$$

$$c = (1 - \widehat{W}_{EE})(1 + \widehat{W}_{II}) + \widehat{W}_{EI}\widehat{W}_{IE}. \tag{S56}$$

The discriminant is

$$\Delta = b^2 - 4ac, \tag{S57}$$

and the two algebraic roots are

$$\mu_\pm(L_B) = \frac{-b \pm \sqrt{\Delta}}{2a}, \qquad q_\pm(L_B) = \sqrt{\mu_\pm(L_B)}. \tag{S58}$$

Equation (**??**) makes the luminance dependence explicit: $L_B$ changes $r_E^{(0)}$ and $r_I^{(0)}$, which changes $s_E$ and $s_I$, which changes the hatted interaction parameters, and therefore changes the roots of the characteristic equation.

Under the $e^{iqx}$ convention, the real part of a bounded complex root is the oscillatory spatial wavenumber, whereas the imaginary part determines spatial attenuation. The square-root signs are therefore selected so that the modes do not grow away from the source, and we define

$$\begin{aligned} k_{n,\pm}(L_B) &= \mathrm{Re}\big[q_\pm(L_B)\big], \\ \lambda_\pm(L_B) &= \mathrm{Im}\big[q_\pm(L_B)\big] \geq 0, \\ f_{n,\pm}(L_B) &= \frac{k_{n,\pm}(L_B)}{2\pi}. \end{aligned} \tag{S59}$$

Only finite branch values are retained.

Both algebraic branches are legitimate candidate paths of the same recurrent-network characteristic equation and are shown in Fig. 1G of the main manuscript. The quantitative psychophysical fits in

Figs. 2–3 use the lower, minus-sign branch,

$$k_n(L_B) \equiv k_{n,-}(L_B), \qquad f_n(L_B) \equiv f_{n,-}(L_B). \tag{S60}$$

The short increase immediately after the transition belongs to the continued minus-sign branch; at larger $L_B$ this branch returns toward lower spatial wavenumbers, whereas the plus-sign branch continues toward higher spatial wavenumbers. This continuation rule is a modelling convention and does not imply that the psychophysical measurements uniquely identify one intrinsic mode. No additional switching rule based on decay rate or mode amplitude is imposed.

The $q$ notation is equivalent to the damped-eigenvalue notation $\Lambda = \lambda + ik_n$ used for the Green's function: the $e^{iqx}$ and $e^{-\Lambda x}$ conventions place the oscillatory and attenuating components in opposite parts of the complex exponent.

##### S4.3 Relation among model wavenumbers and experimental spatial frequency

The relevant quantities have distinct definitions:

$$\boxed{\begin{aligned} k_n &: \text{ intrinsic network spatial wavenumber,} \\ k_r &: \text{ resonance spatial wavenumber under periodic forcing,} \\ f_n = k_n/(2\pi) &: \text{ intrinsic ordinary spatial frequency,} \\ f_r = k_r/(2\pi) &: \text{ resonance ordinary spatial frequency,} \\ f_{\text{pref}} &: \text{ experimental preferred spatial frequency.} \end{aligned}} \tag{S61}$$

The model predicts that the periodically forced resonance $k_r$ is shaped by the intrinsic spatial organization characterized by $k_n$. For comparison with the psychophysical measurements, both model wavenumbers are converted to ordinary spatial frequencies. Thus, the central comparison is among $f_{\text{pref}}$, $f_r$, and $f_n$, all expressed in cycles per degree. The $f_n(L_B)$ and $f_r(L_B)$ curves shown in the main manuscript were fitted independently rather than constrained to be numerically identical.

#### S5 Preferred spatial frequency from psychophysical measurements

##### S5.1 General definition

Let $\widetilde{S}(f)$ denote a smooth fitted representation of a measured contrast-sensitivity function. The definition of preferred spatial frequency does not depend on one particular fitting family: another suitable smooth CSF can be substituted without changing the definition. The preferred spatial frequency is the frequency at which the fitted CSF attains its maximum. At a smooth interior maximum,

$$\left.\frac{d\widetilde{S}(f)}{df}\right|_{f=f_{\text{pref}}} = 0, \qquad \left.\frac{d^2\widetilde{S}(f)}{df^2}\right|_{f=f_{\text{pref}}} < 0. \tag{S62}$$

#### S5.2 Phenomenological CSF used in the present analysis

For the present data, contrast-sensitivity estimates were interpolated using the phenomenological exponential-plus-Gaussian function

$$S(f) = b \exp\left[-\left(\frac{f - f_0}{f_2}\right)\right] + a \exp\left[-\left(\frac{f - f_0}{f_1}\right)^2\right], \qquad f_2 = 1. \tag{S63}$$

The first exponential term is intentionally neither squared nor expressed using an absolute value; Eq. (S58) reproduces the function implemented in the fitting code. The function is used only as a smooth phenomenological interpolant and its individual parameters are not assigned physiological meaning. This treatment is motivated by established analytical descriptions of the CSF [Watson and Ahumada, 2005, Campbell et al., 1966, Watson, 2006].

#### S5.3 Resampling estimation of preferred spatial frequency

Each adaptive staircase was terminated after 10 reversals. To propagate uncertainty in the staircase threshold through to the preferred-spatial- frequency estimate, the reversal data were resampled. On each resampling iteration, eight of the 10 reversals obtained for each completed staircase were randomly selected *without replacement* and used to estimate the staircase contrast threshold. Threshold estimates from the available staircases were then combined for each spatial-frequency condition and converted to contrast sensitivity, yielding one resampled CSF. Unlike the primary point estimator, which used the final eight reversals (reversals 3–10), this resampling pool included all 10 completed reversals, including reversals 1 and 2.

For a single staircase there are

$$\binom{10}{8} = 45 \tag{S64}$$

possible eight-reversal subsets. A complete CSF resample combines such subsets across all completed staircases and spatial-frequency conditions. Consequently, although each individual staircase has only 45 possible eight-reversal subsets, the number of possible complete resampled CSFs is much larger because selections are combined across conditions.

For each subject and luminance condition, 10,000 resampling iterations were performed. The phenomenological CSF in Eq. (S58) was fitted independently to every resampled CSF, and the preferred spatial frequency $f_{\text{pref}}$ was extracted as the spatial frequency at which the fitted function reached its maximum. Thus, each subject and luminance condition produced an initial distribution of 10,000 candidate $f_{\text{pref}}$ estimates.

Fit quality was assessed using the normalized root-mean-square error (NRMSE). If $S_i$ is the measured contrast sensitivity at frequency $f_i$ and $\widehat{S}(f_i)$ is the corresponding fitted value, the quantity used in the analysis was

$$\text{NRMSE} = \frac{\sqrt{N^{-1} \sum_{i=1}^{N} \left[S_i - \widehat{S}(f_i)\right]^2}}{N^{-1} \sum_{i=1}^{N} S_i}, \tag{S65}$$

where $N$ is the number of sampled spatial frequencies in that CSF.

Fits satisfying the global criterion

$$0 \leq \text{NRMSE} \leq 0.3 \tag{S66}$$

were retained. The upper bound of 0.3 was chosen as a permissive fit-quality criterion: fits with larger NRMSE values typically provided visibly unsuitable descriptions of the resampled CSFs, whereas the chosen bound removed these poor fits while retaining substantial variability among otherwise plausible fits.

Preferred-spatial-frequency estimates at or below 1.05 cycles/degree were subsequently excluded. This lower cutoff was introduced because poorly constrained fits could occasionally produce spurious low-frequency maxima or secondary modes. It therefore restricted the summary statistics to the range considered reliable for estimation of the CSF peak and served as a fit-quality safeguard rather than as a constraint on the luminance dependence of the model. No analogous upper-frequency exclusion criterion was applied.

The reported preferred spatial frequency was the mean of the retained $f_{\text{pref}}$ distribution,

$$\overline{f}_{\text{pref}} = \frac{1}{N_{\text{ret}}} \sum_{m=1}^{N_{\text{ret}}} f_{\text{pref},m}, \tag{S67}$$

where $N_{\text{ret}}$ is the number of resampled fits remaining after the fit-quality and lower-frequency criteria were applied. Error bars were defined by the 2.5th and 97.5th percentiles of this retained distribution, giving a 95% percentile interval. The resampling procedure therefore propagates uncertainty from the staircase reversals through threshold estimation, CSF fitting, and finally the inferred peak location $f_{\text{pref}}$. Because the procedure perturbs the observed reversal set rather than resampling independent observations from an underlying population, this percentile range measures within-dataset estimator stability and is not interpreted as a classical non-parametric bootstrap confidence interval.

#### S6 Mapping physical luminance to model background drive

The mathematical derivations above use $L_B$ for the uniform background drive of the network. During fits to the psychophysical measurements, the experimentally measured luminance is kept distinct from this model variable. Let $L_v$ denote physical luminance in cd/m$^2$. The measured luminance was mapped to the model drive according to

$$X(L_v) = \log_{10}\left(\frac{L_v + L_{\text{off}}}{L_{\text{ref}}}\right), \tag{S68}$$

with

$$L_{\text{off}} = 100\ \text{cd/m}^2, \qquad L_{\text{ref}} = 1\ \text{cd/m}^2. \tag{S69}$$

The reference luminance makes the logarithm dimensionless. The offset and the multiplicative scale factor (fixed at 1) were not optimized during fitting.

The transformed input was divided between the populations as

$$i_E^{(0)} = \alpha X, \qquad i_I^{(0)} = (1 - \alpha)X. \tag{S70}$$

Thus, when model curves are evaluated against the psychophysical data, the uniform background drive appearing in the model equations is supplied by the transformed quantity $X(L_v)$. The logarithmic horizontal axes in the main figures display the original physical luminance values and are only a plotting convention; they do not constitute an additional transformation of the model input.

### S7 Fitting procedures

#### S7.1 Empirical CSF fits

Unless otherwise stated, nonlinear least-squares optimization was performed using the Trust Region Reflective algorithm implemented by `scipy.optimize.curve_fit`. For the empirical CSFs, no constraints were imposed on the interpolation parameters during optimization. The same fitting procedure was applied independently to each resampled CSF described above. The fitted parameters themselves were not used as mechanistic estimates; only the location of the fitted maximum was propagated into the preferred-spatial-frequency analysis.

#### S7.2 Fits of intrinsic spatial frequency to psychophysical data

For each candidate parameter set and each measured luminance $L_v$:

(1) The physical luminance was mapped to the model input $X(L_v)$ using Eq. (S62).

(2) The homogeneous background state was determined from Eqs. (S13)–(S14), with the transformed drive supplied to the background-input terms.

(3) The local gains $s_E$ and $s_I$ and all hatted interaction parameters were recomputed for that background state.

(4) The minus-sign root was evaluated as $k_{n,-}(L_B) = \mathrm{Re}[q_-(L_B)]$ and converted to ordinary spatial frequency using $f_n(L_B) = k_{n,-}(L_B)/(2\pi)$.

(5) Nonlinear least squares was used to minimize the discrepancy between the predicted $f_n$ curve and the measured preferred spatial frequencies.

The recurrent and lateral coupling strengths, the input partition $\alpha$, and the background-state parameters used in the numerical implementation were included in the optimization described in the main manuscript. The resulting parameter set was used to evaluate $f_n$ across the measured luminance range.

#### S7.3 Fits of resonance spatial frequency to psychophysical data

The resonance curve was fitted independently from the intrinsic-frequency curve. For each candidate parameter set and luminance level:

(1) The background state and renormalized parameters were recomputed.

(2) Trial ordinary spatial frequencies $f$ were converted to wavenumbers using $k = 2\pi f$, and the signed excitatory response coefficient $\xi(2\pi f, L_B)$ was evaluated using Eq. (S39).

(3) The frequency at which $\xi$ reached its maximum was recorded as $f_r$; no absolute value was applied.

(4) Nonlinear least squares was used to fit the resulting $f_r$ values to the measured preferred spatial frequencies.

The $f_n$ and $f_r$ curves therefore provide separate tests of whether the intrinsic and periodically forced descriptions of the recurrent network can reproduce the luminance dependence of the psychophysical preferred spatial frequency.

##### S7.4 Fits to published physiological data

The recurrent E–I framework was also fitted to the previously published physiological measurements summarized in Fig. 4 of the main text.

**Figure 4A.** Seven of the eight coupling weights were held fixed, while one coupling weight was varied between the experimental stimulus conditions. This analysis tested whether a change in a single network interaction could reproduce the family of normalization-like cortical response curves.

**Figure 4B.** The model was fitted separately to the population-averaged cortical hyperpolarization measurements. The recurrent network was numerically integrated to steady state for each luminance-step condition, after which scaling and offset parameters mapped the model response onto the normalized experimental measurements.

**Figure 4C.** The excitatory and inhibitory responses were fitted simultaneously to the published excitatory postsynaptic current (EPSC) and inhibitory postsynaptic current (IPSC) measurements. The recurrent E–I model was numerically integrated to steady state for each luminance condition. Shared recurrent coupling parameters were estimated by minimizing the combined residual error across the excitatory and inhibitory datasets. Linear scaling and offset parameters mapped the model responses onto the normalized experimental measurements.

These fits are intended to assess whether a common recurrent E–I architecture can reproduce the qualitative response structure of the published physiological data. They should not be interpreted as unique estimates of biological synaptic parameters.

#### S8 Computational sequence and interpretation

For clarity, the complete luminance-dependent calculation can be summarized as the following sequence:

$$\begin{gathered} L_v \longrightarrow X \longrightarrow \left(r_E^{(0)}, r_I^{(0)}\right) \longrightarrow (s_E, s_I) \longrightarrow \left(\widehat{W}, \widehat{D}\right), \\ (\widehat{W}, \widehat{D}) \longrightarrow \begin{cases} q_-(L_B) \longrightarrow f_n, & \text{intrinsic spatial-mode calculation,} \\ \xi(2\pi f, L_B) \longrightarrow f_r, & \text{periodically forced calculation.} \end{cases} \end{gathered} \tag{S71}$$

The psychophysical estimate is obtained through a separate resampling pipeline,

$$\begin{aligned}\text{staircase reversals} &\longrightarrow \text{eight-of-ten resampling} \longrightarrow \text{threshold estimates}\\ &\longrightarrow \text{resampled CSF} \longrightarrow \widetilde{S}(f) \longrightarrow f_{\text{pref}}\\ &\longrightarrow \text{fit-quality screening} \longrightarrow \left(\overline{f}_{\text{pref}},\ 95\%\ \text{percentile interval}\right).\end{aligned} \tag{S72}$$

The central comparison in the paper is therefore between the perceptual peak location $f_{\text{pref}}$ and the ordinary spatial frequencies $f_n$ and $f_r$ predicted by the recurrent network, rather than between the individual parameters of the empirical CSF fit and the biological coupling parameters.

#### S9 Scope of the linear analysis

The supplementary derivation is a first-order analysis about a luminance-dependent homogeneous operating state. Several consequences follow.

First, the anatomical coupling weights $w_{PQ}$ and $\widetilde{w}_{PQ}$ are not themselves changed by luminance. Luminance changes $r_E^{(0)}$ and $r_I^{(0)}$, which changes the local slopes $s_E$ and $s_I$ of the nonlinear response functions and thereby changes the renormalized parameters $\widehat{W}_{PQ}$ and $\widehat{D}_{PQ}$.

Second, the periodic response and the intrinsic response are distinct calculations. The resonance wavenumber $k_r$ depends on the response to an external periodic stimulus, whereas $k_n$ characterizes the spatial organization of the recurrent operator without that periodic forcing. Their ordinary spatial-frequency representations are $f_r$ and $f_n$.

Third, the experimentally defined $f_{\text{pref}}$ is not imposed on the model. It is estimated from the measured CSF and subsequently compared with $f_r$ and $f_n$.

Finally, the present analysis concerns the luminance-dependent spatial organization of the model around each background operating state. It does not require luminance to alter the underlying anatomical connectivity.
